# pHluo_M153R-CD63, a bright, versatile live cell reporter of exosome secretion and uptake, reveals pathfinding behavior of migrating cells

**DOI:** 10.1101/577346

**Authors:** Bong Hwan Sung, Ariana von Lersner, Jorge Guerrero, David Inman, Roxanne Pelletier, Andries Zijlstra, Suzanne M. Ponik, Alissa M. Weaver

## Abstract

Small extracellular vesicles called exosomes affect a variety of autocrine and paracrine cellular phenotypes, including cellular migration, immune activation, and neuronal function. Understanding the function of exosomes requires a variety of tools, including live cell imaging. We previously constructed a live-cell reporter, pHluorin-CD63, that allows dynamic subcellular monitoring of exosome secretion in migrating and spreading cells. However, there were some caveats to its use, including dim fluorescence and the inability to make cell lines that stably express the protein. By incorporating a stabilizing mutation in the pHluorin moiety, M153R, pHluorin-CD63 now exhibits higher and stable expression in cells and superior monitoring of exosome secretion. Using this improved construct, we demonstrate visualization of secreted exosomes in 3D culture and *in vivo* and identify a role for exosomes in promoting leader-follower behavior in 2D and 3D migration. By incorporating a further non-pH-sensitive red fluorescent tag, this reporter allows visualization of the exosome lifecycle, including multivesicular body (MVB) trafficking, MVB fusion, exosome uptake and endosome acidification. This new reporter will be a useful tool for understanding both autocrine and paracrine roles of exosomes.

## Introduction

Extracellular vesicles (EVs) are nano-sized vesicles released from cells with potent autocrine and paracrine biological activities^1^. Although EVs were first considered to be cell debris with little biological relevance, EVs are now understood to constitute a fundamental mode of cell-cell communication that mediates delivery of specific protein, nucleic acid and lipid cargoes to recipient cells in diverse contexts^1, 2^. Indeed, EVs are involved in the pathogenesis of diverse diseases, including infections^3^, neurodegenerative disorders^4^, cardiovascular disease^5^, and cancer^6, 7^.

EVs can be classified by their size, biogenesis mechanism, cargoes, or density, e.g. small EVs, including exosomes, and larger EVs such as shed microvesicles (MVs) and large oncosomes^8^. Exosomes are a type of small EV with diameter of 30 to 150 nm and have been the most studied with respect to their involvement in EV function^1, 2^. Exosomes are formed as intraluminal vesicles (ILVs) in late endosomal organelles called multivesicular bodies (MVBs) and secreted after fusion of MVBs with the plasma membrane. Several exosome biogenesis processes have been proposed, including capture of ubiquitinated cargoes by the endosomal sorting complex required for transport (ESCRT) machinery ^9^. Syndecan-syntenin complexes also regulate exosome biogenesis via the ESCRT accessory protein Alix ^10^. An ESCRT-independent biogenesis mechanism involving membrane curvature induced by ceramide generation by neutral sphingomyelinase 2 (nSMase2) has also been described^11^. Tetraspanin proteins such as CD9, CD63, and CD81 are frequently used markers of exosomes and other small EVs and may be involved in ILV cargo selection and/or biogenesis ^12^.

CD63 is a member of the tetraspanin superfamily, is enriched on ILVs in late endosomal MVBs, and is a broadly-used classic exosomal marker. Knockout expression of CD63 by CRISPR/Cas9 impairs secretion of small EVs but not LEVs, suggesting that it contributes to exosome biogenesis ^13^. CD63 has been used to label and track exosomes and MVBs in many studies^14-18^. However, most previous studies used pH-insensitive fluorescent proteins such as GFP or RFP, leading to extremely bright fluorescence of internal endosomes. This bright internal fluorescence limits the ability to observe fusion events of MVBs with the plasma membrane due to a poor signal-to-noise ratio. To solve this problem, we adapted an approach from the synaptic vesicle field, that leverages the properties of a pH-sensitive GFP derivative, pHluorin, to observe dynamic vesicle fusion events ^19^. pHluorin is virtually non-fluorescent under acidic conditions but fluoresces at neutral pH ^19^. We first developed a pHluorin-tagged CD63 reporter to track exosome secretion and used it to demonstrate that MVB fusion precedes adhesion formation in spreading cells by 1-2 min ^20^. We also observed that pHluorin-CD63-positive adhesive trails were left behind migrating cells ^20^. Subsequently, a similar reporter, with the pHluorin group placed 7 amino acids away from ours in the first extracellular loop of CD63, was used to study GPCR regulation of exosome secretion ^21^.

pHluorin-CD63 is a powerful tool to track exosome secretion and MVB fusion with the plasma membrane. However, while this construct is useful in 2-dimensional (2D) tissue culture conditions with high resolution imaging, the pHluorin moiety is subject to degradation in cells^22^, leading to low levels of expression and fast photobleaching. Thus, we have found it to be difficult to stably express it in cells, which limits its use for live imaging to select conditions.

In this study, we improved the stability and brightness of superecliptic pHluorin-CD63 by incorporating a single amino acid mutation, M153R, previously identified to stabilize ratiometric pHluorin in bacterial fusions ^22^. We demonstrate that the mutated pHluorin-CD63, pHluo_M153R-CD63, can now be expressed as a stable construct in cells and is a bright reporter for exosome secretion. Using this construct, we are able to resolve individual exosome puncta and observe pathfinding behavior of migrating cells along extracellularly deposited exosome trails in both 2D and 3D. By tagging an additional pH-insensitive red fluorescent protein to pHluo_M153R-CD63, we are further able to track multiple aspects of the exosome lifecycle, including MVB movements within cells before fusion, endocytosis of extracellular exosome deposits, and acidification of exosome-containing endocytic compartments.

## Results

### Mutation of pHluorin-CD63 creates a bright, stable live imaging reporter

To improve on our previous reporter, we tested whether a mutation previously shown to stabilize ratiometric pHluorin in bacterial fusion proteins, M153R^22^, would also stabilize our superecliptic pHluorin-CD63 construct. The pHluorin-CD63 construct from our previous study^20^ was mutated on Methionine 153 to Arginine by site-directed mutagenesis (pHluo_M153R-CD63, Fig. 1a). As shown in Fig 1b, CD63 is a tetraspanin protein with two extracellular loops. The pHluorin is inserted into the small extracellular loop at position 43. Upon fusion of MVB with the plasma membrane, the pHluorin moiety is exposed to neutral pH and becomes fluorescent, which enables dynamic monitoring of exosome secretion (Fig. 1b). After site-directed mutagenesis, pHluo_M153R-CD63 was cloned into a lentiviral vector and stably expressed in HT1080 fibrosarcoma cells. To test whether pHluo_M153R-CD63 labels small EVs, conditioned media of pHluo_M153R-CD63-expressing HT1080 cells were collected and serially centrifuged. Large EVs, typically consisting of shed microvesicles, were pelleted through a 10,000 × g spin for 30 min and small EVs, typically containing exosomes, were pelleted by centrifugation at 100,000 × g overnight. Nanoparticle tracking analysis (NTA) of small EVs showed the expected size distribution for exosomes with a peak diameter of 105 nm whereas large EVs had peak diameters of 195 nm and 405 nm (Fig. 1c). Consistent with the previously reported role of pHluorin-CD63 as a reporter of MVB fusion and exosome secretion, immunoblotting of cell lysates and purified EVs revealed that pHluo_M153R-CD63 is exclusively detected in the exosome-enriched small EV preparation, and not in the larger EVs (Fig. 1d). NTA showed an increased secretion rate of small EVs from pHluo_M153R-CD63-expressing cells compared with parental HT1080s but no change in the number of large EVs (Supplementary Fig. 1a). Live imaging of HT1080 cells stably expressing pHluo_M153R-CD63 as well as the plasma membrane marker mCherry-CaaX revealed numerous pHluo_M153R-CD63-positive puncta left behind migrating HT1080 cells. These puncta were mCherry-CaaX-negative, suggesting that the deposits are likely to be exosomes and not plasma membrane-derived MVs or debris (Fig. 1e and Supplementary Video 1, upper panel). These findings are similar to the previous green fluorescent “slime trails”, that we observed left behind cells transiently transfected with pHluorin-CD63 (^20^, Fig 1f); however, the deposited trails were much brighter and more easily resolved into puncta using standard epifluorescence imaging (Fig. 1e and Supplementary Video 1. Note that pHluo-CD63 fluorescence in f and the lower panel of the movie is much dimmer). Also, Western blots of lysates from cells transiently transfected with either pHluorin-CD63^20^ or pHluorin_M153R-CD63 revealed that our previous construct is present at lower levels than pHluorin_M153R. These data suggest that pHluo_M153R-CD63 is indeed more stable (Supplementary Fig. 1b, arrows). Consistent with that idea, we find that the new reporter can be stably expressed in cells using lentiviral transduction, which has many advantages, including the ability to FACS sort populations for more uniform fluorescent expression and to image in many more conditions, potentially including low light, lower resolution, 3D and *in vivo*.

**Figure 1.**
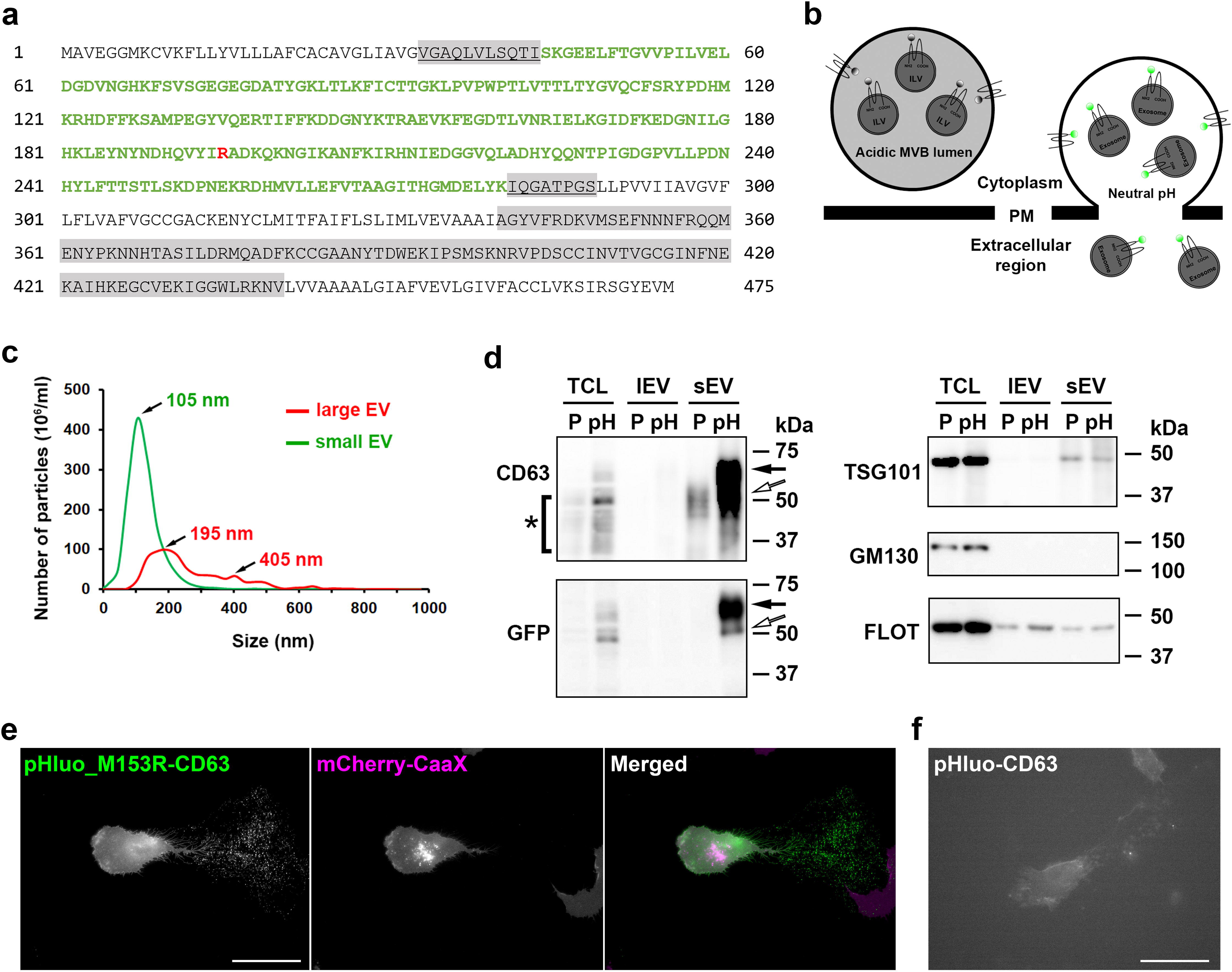
pHluorin_M153R-CD63 is a bright, stable exosome reporter. (**a**) Sequence of pHluorin_M153R-CD63. pHluorin sequence is in green color. Highlighted regions in grey represent small (underlined) and large extracellular loops. M153R mutation is marked in red. (**b**) Diagram of pHluorin_M153R-CD63 construct. Notice pHluorin_M153R tag has bright fluorescence upon fusion of the multivesicular body (MVB) with the plasma membrane due to the exposure to neutral pH. Otherwise, it is non-fluorescent in the acidic condition of the MVB lumen. ILV, intraluminal vesicle. PM, plasma membrane. (**c**) Representative trace from nanoparticle tracking analysis of large EVs (lEVs) and small EVs (sEVs). (**d**) Western blot analysis of cells and EVs with anti-CD63, anti-GFP, EV markers (TSG101, Flotillin) and Golgi marker (GM130). TCL, total cell lysate. lEV, large EV. sEV, small EV. P, parental cells. pH, pHluorin_M153R-CD63-expressing cells. Black arrows indicate full length pHluorin_M153R-tagged CD63, which is shifted due to the GFP moiety of 27 kDa, while white arrows indicate potential cleaved form of CD63 tagged with pHluorin_M153R. Asterisk indicates cellular CD63, which has a broad range due to glycosylation. (**e**) Still images from Supplementary Video 1 (upper panel) showing a migrating HT1080 cell stably expressing mCherry-CaaX (magenta) and pHluo_M153R-CD63 (green). Colocalization of magenta and green is white. Notice that the deposits left behind the migrating cell are only labeled with CD63 not with CaaX. Scale bar, 50 µm. Movie representative of 38 movies. (*f*) Still images from Supplementary Video 1 (lower panel) showing a migrating HT1080 cell transiently expressing pHluo-CD63. Scale bar, 50 µm. Movie representative of 12 movies.

### Extracellular pHluo_M153R-CD63 puncta correspond to exosome deposits

In a previous study, correlative light-transmission electron microscopy imaging revealed that fluorescent flashes of pHluorin-CD63 at the plasma membrane indeed correspond to MVB fusion events ^21^. To determine whether pHluo_M153R-CD63 likewise reports exosome secretion, we knocked down the MVB docking protein Rab27a^23^ with shRNA in pHluo_M153R-CD63-expressing HT1080 cells (Fig. 2a). As expected, the number of small EVs released into the media of Rab27a-KD cells was greatly decreased compared to control cells, as assessed by NTA (Fig. 2b). While not all small EVs are expected to be exosomes^24-26^, a substantial portion of them are expected to derive from MVBs. Imaging of control and Rab27a-KD cells expressing pHluo_M153R-CD63 revealed greatly reduced extracellular deposition of pHluo_M153R-CD63-positive puncta by Rab27a-KD cells (Fig. 2c), as quantitated by measuring the area and integrated intensity of the pHluo_M153R-CD63-positive deposits surrounding the cells (Fig. 2d and e). pHluo_M153R-CD63-positive deposition also was observed from other cell types (Supplementary Fig. 2). Some of the puncta are brighter than others, suggesting that they may represent groups of exosomes. Furthermore, some puncta are arranged in linear trails, which may represent organization by cells as they migrate over them (e.g. as in Supplementary Video 1, trailing edge of migrating cell).

**Figure 2.**
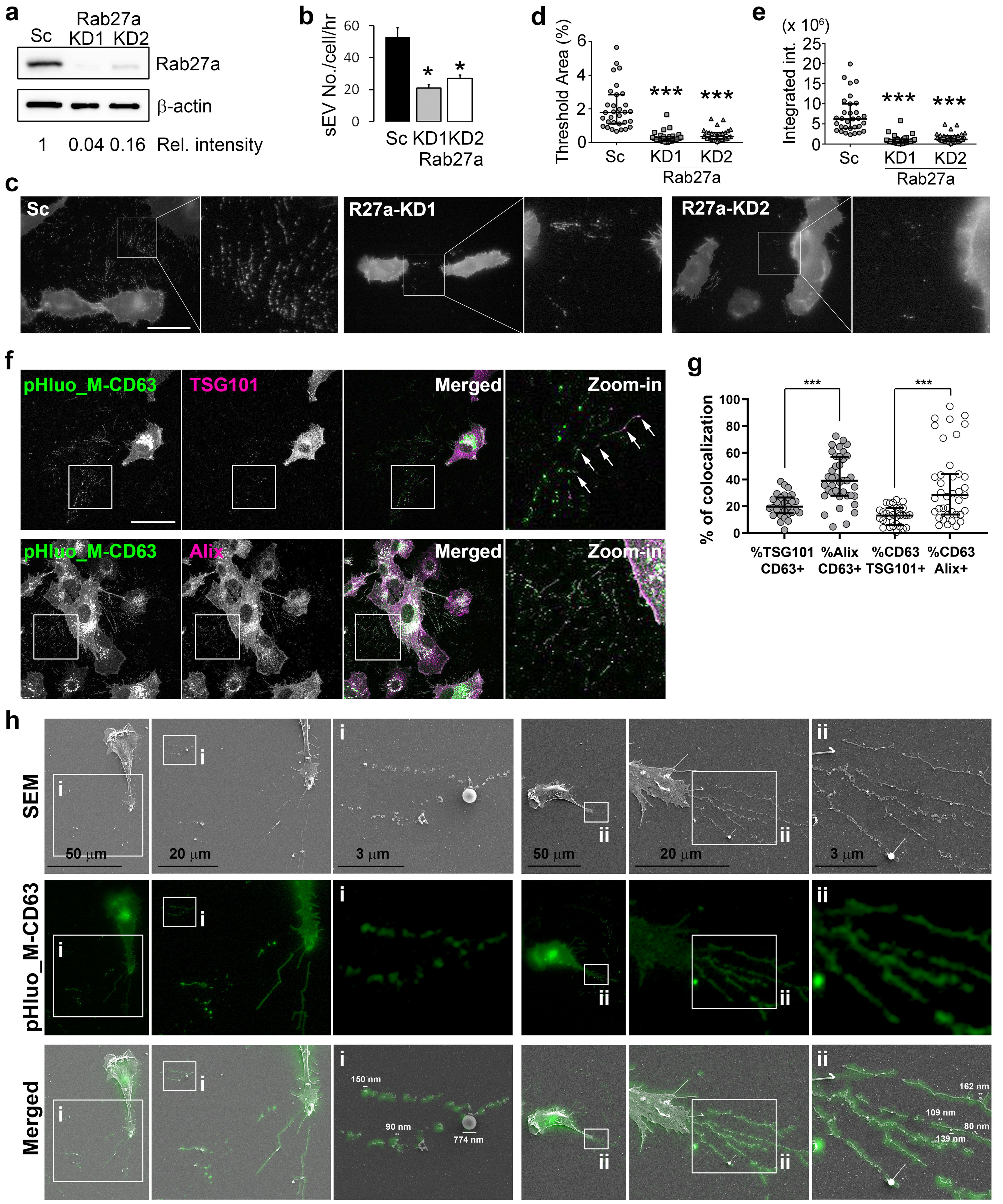
pHluorin_M153R-CD63-positive trails mark secreted exosomes. (**a**) Immunoblotting of Rab27a in pHluo_M153R-CD63-stably expressing HT1080. Sc, scrambled control. KD, knockdown. (**b**) Comparison of sEV secretion rate between control and Rab27a-KDs from 3 independent experiments. (**c-e**) Live imaging and analysis of extracellular pHluorin_M153R-CD63 for control and Rab27a-KD cells. (**c**) Representative images, n=31 from 3 independent experiments for each cell line. Scale bar, 30 µm. (**d**) Threshold area of the extracellular region from (**c**) shown as a scatter-plot with median and quartile range. (**e**) Integrated Intensity of the extracellular region from (**c**) shown as a scatter-plot with median and quartile range. (**f**) Representative images of immunofluorescent staining of fixed cells for the exosomal markers TSG101 or Alix (magenta) and pHluorin (green). Arrows indicate colocalization of TSG101 (magenta) with pHluo_M153R-CD63-positive puncta (green) to form white. Note many white puncta in the CD63+Alix merged images. Scale bar, 30 µm. (**g**) Colocalization analysis from 29 and 39 images for TSG101 and Alix, respectively, shown as a scatter-plot with median and quartile range. (**h**) Correlative light-electron microscopy of pHluo_M153R-CD63-stably expressing HT1080. Increasing magnification of images are shown to the right and indicated with i or ii. Representative of 22 cells from 4 experiments. \**P*<0.05; \*\*\**P*<0.001.

As further confirmation that the extracellular pHluo_M153R-CD63-positive puncta are exosomes, we colocalized them with additional exosome markers, specifically those associated with the ESCRT machinery that causes intraluminal vesiculation within MVB (Fig. 2f). Immunostaining of fixed cells revealed that the ESCRT-I protein TSG101 was present in ∼20% of extracellular pHluo_M153R-CD63 puncta. Immunostaining with the ESCRT accessory protein, Alix, revealed even more colocalization with pHluo_M153R-CD63-positive extracellular puncta (median ∼40%, Fig. 2g). Analysis of the percent TSG101 or Alix that were positive for CD63 likewise revealed more colocalization with Alix than TSG101, in some cells almost 100% colocalization. The difference in colocalization with TSG101 versus Alix is consistent with the well-known heterogeneity of exosomes as well as with previous immunofractionation experiments in which an exosome biogenesis syndecan-syntenin-Alix complex was present in EVs immunoprecipitated with anti-CD63 antibody^10^. Of note, due to neutralization of intracellular pH by the paraformaldehyde fixation procedure, pHluo_M153R-CD63 shows bright fluorescence in internal endosomal structures in these images (Fig. 2f).

To further determine whether pHluo_M153R-CD63 extracellular puncta and trails represent exosome deposition, we performed correlative light-electron microscopy in which HT1080 cells expressing pHluo_M153R-CD63 were first observed by epifluorescence microscopy followed by scanning electron microscopy (Fig. 2h). pHluo_M153R-CD63 fluorescence corresponded to small EVs with diameters ranging from 80 to 160 nm, which were frequently organized into linear trails (Fig 2h-i) and/or associated with retraction fibers (Fig. 2h-ii). By contrast, larger vesicles (e.g. the 774 nm EV in Fig 2h-i) were occasionally observed and were not fluorescent. These data are consistent with CD63 labeling primarily exosomes and not larger microvesicles^20, 27^. We also observed rare fluorescent pouches of small vesicles left behind cells (Supplementary Fig. 3). These pouches resemble previously described migrasomes^28^, which are reportedly driven by a different tetraspanin, TSPAN4, or could alternatively represent groups of exosomes that derived from MVB fusion and tore off plasma membrane during cell migration. Additional mechanistic experiments would be required to identify their origin.

**Figure 3.**
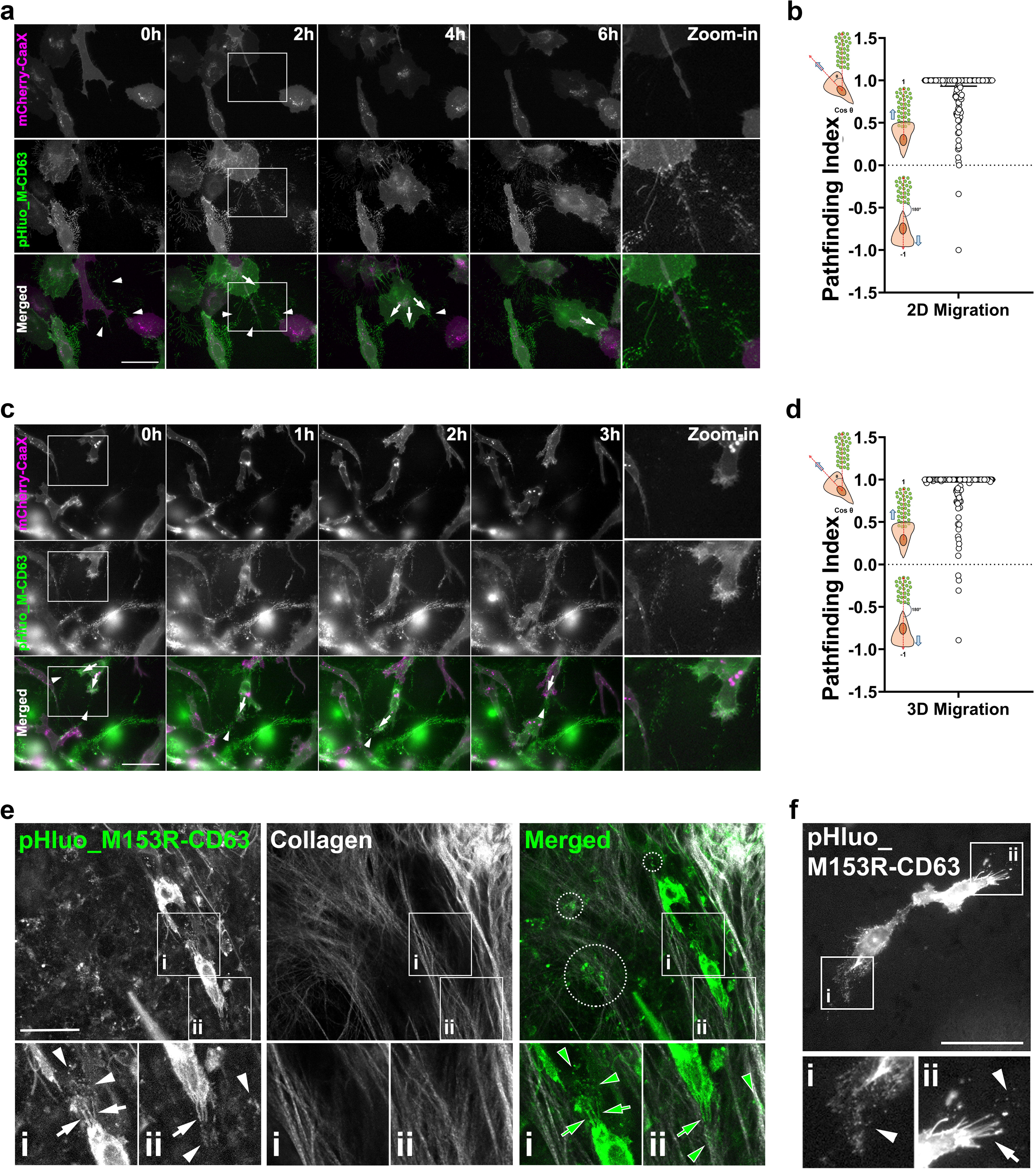
Cells exhibit pathfinding behavior on exosome trails. (**a**) Time series images from live imaging of pHluorin_M153R-CD63/mCherry-CaaX dual-expressing HT1080 cells in 2D culture conditions (see Supplementary Video 2). mCherry-CaaX is shown in magenta and pHluorin_M153R-CD63 is shown in green in the merged image. n=186 cells from 31 movies from 4 independent experiments. Arrowheads indicate exosome trails. Arrows indicate protrusions. (**b**) Pathfinding index from (**a**) shown as a scatter-plot with median and quartile range. (**c**) Time series images from live imaging of pHluorin_M153R-CD63/mCherry-CaaX dual-expressing HT1080 cells in 3D collagen gels (see Supplementary Video 3). n=151 cells from 27 movies from 4 independent experiments. Arrowheads indicate exosome trails. Arrows indicate protrusions. (**d**) Pathfinding index from (**c**) shown as a scatter-plot with median and quartile range. (**e**) Representative intravital images of pHluo_M153R-CD63 in mice. Arrows indicate adhesive protrusions. Arrowheads indicate exosome deposits. Examples of exosome deposits associated with collagen fibers are shown in dotted circles. (**f**) Representative intravital images of pHluo_M153R-CD63 in chicks. Arrows indicate adhesive protrusions. Arrowheads indicate exosome deposits. Scale bars, 50 µm.

### Cells exhibit pathfinding behavior over pHluo_M153R-CD63 deposits

Previous reports indicate that exosome secretion promotes directional migration of several cell types including cancer cells^20, 29^, neutrophils^30^ and Dictyostelia^31^. To visualize dynamically how secreted exosomes influence cell migration, we performed live imaging under diverse conditions. In 2-dimensions, mCherry-CaaX/pHluo_M153R-CD63-double labeled HT1080 cells exhibited path-finding behavior along exosomes. Thus, cells extended leading edge protrusions toward and over exosome deposits labeled with pHluo_M153R-CD63 (Fig. 3a, arrows, Supplementary Video. 2) and then migrated along the deposits (Fig. 3a, arrow heads). We observed the same phenotype in 3D collagen gels (Fig. 3c, Supplementary Video. 3). To provide a quantitative assessment of this behavior, we analyzed the relationship of cell migration paths relative to exosomes by measuring the angle of each cell trajectory to the nearest exosome trail. We then took the cosine of the angle to obtain the pathfinding index. Thus, zero deviation of the path from an exosome trail is 1, i.e. if cells migrated directly toward the nearest exosome deposits or if cells migrated directly over the deposits. If cells migrated away from the nearest deposits in the opposite direction with a 180° angle, then the pathfinding index would be −1, as shown in the cartoons in Fig 3b and d. For both 2D and 3D migration, HT1080 cells primarily migrated toward or over exosome deposits with a median pathfinding index of 1 and 0.0607 and 0.0254 quartile ranges for 2D and 3D migration, respectively (Fig 3b and d).

To further examine whether pHluo_M153R-CD63 extracellular puncta are visible in living systems, we performed intravital imaging in mice as well as in chick embryos. For the mouse system, MDA-MB-231 breast cancer cells stably expressing pHluo_M153R-CD63 were injected into the 4^th^ mammary fat pad and observed 7-10 days later by multiphoton imaging through a mammary imaging window. For the chick embryo, HT1080 fibrosarcoma cells expressing pHluo_M153R-CD63/mCherry-CaaX were injected into the chorioallantoic vein and imaged after 24 hours using an upright epifluorescence microscope. In both systems, bright fluorescent pHluo_M153R-CD63-positive puncta were observed surrounding the pHluo_M153R-CD63-expressing cancer cells (Fig. 3e and f, arrowheads). Similar to our *in vitro* observations, these puncta were frequently associated with long cellular extensions that could be either retraction fibers or filopodia (Fig. 3e and f, arrows). In the mouse xenograft system, some exosome deposits appeared to adhere to collagen fibers, as observed by second-harmonic generation microscopy (Fig. 3e, dotted circles). These extracellular exosome deposits could theoretically provide a substrate for cell migration as we previously proposed ^20^, and now observe in 2D and 3D (Fig 3a-d).

### Creation of a dual color reporter for MVB trafficking and fusion, and exosome uptake

We previously observed by TIRF microscopy that putative MVB fusion events closely precede adhesion formation^20^ in spreading cells. However, our ability to track and observe fusion events in other contexts (e.g. by widefield or confocal microscopy) was limited due to the dim fluorescence of the original pHluorin-CD63 construct, and lack of a pH-insensitive tag to track MVB trafficking events before fusion with the plasma membrane. To solve the latter problem, mScarlet, a bright monomeric red fluorescent protein^32^ was cloned at the C-terminus of pHluo_M153R-CD63 to make pHluo_M153R-CD63-mScarlet. This construct exhibits red fluorescence under acidic conditions (e.g. when contained in the MVB lumen) and dual fluorescence under neutral conditions (e.g. after secretion). Live confocal microscopy of HT1080 cells stably expressing pHluo_M153R-CD63-mScarlet showed that MVBs are frequently trafficked to protruding cell edges and fuse there (Fig. 4a and Supplementary Video 4). Note the frequent appearance of pHluo_M153R-CD63-mScarlet, often before membrane protrusion.

**Figure 4.**
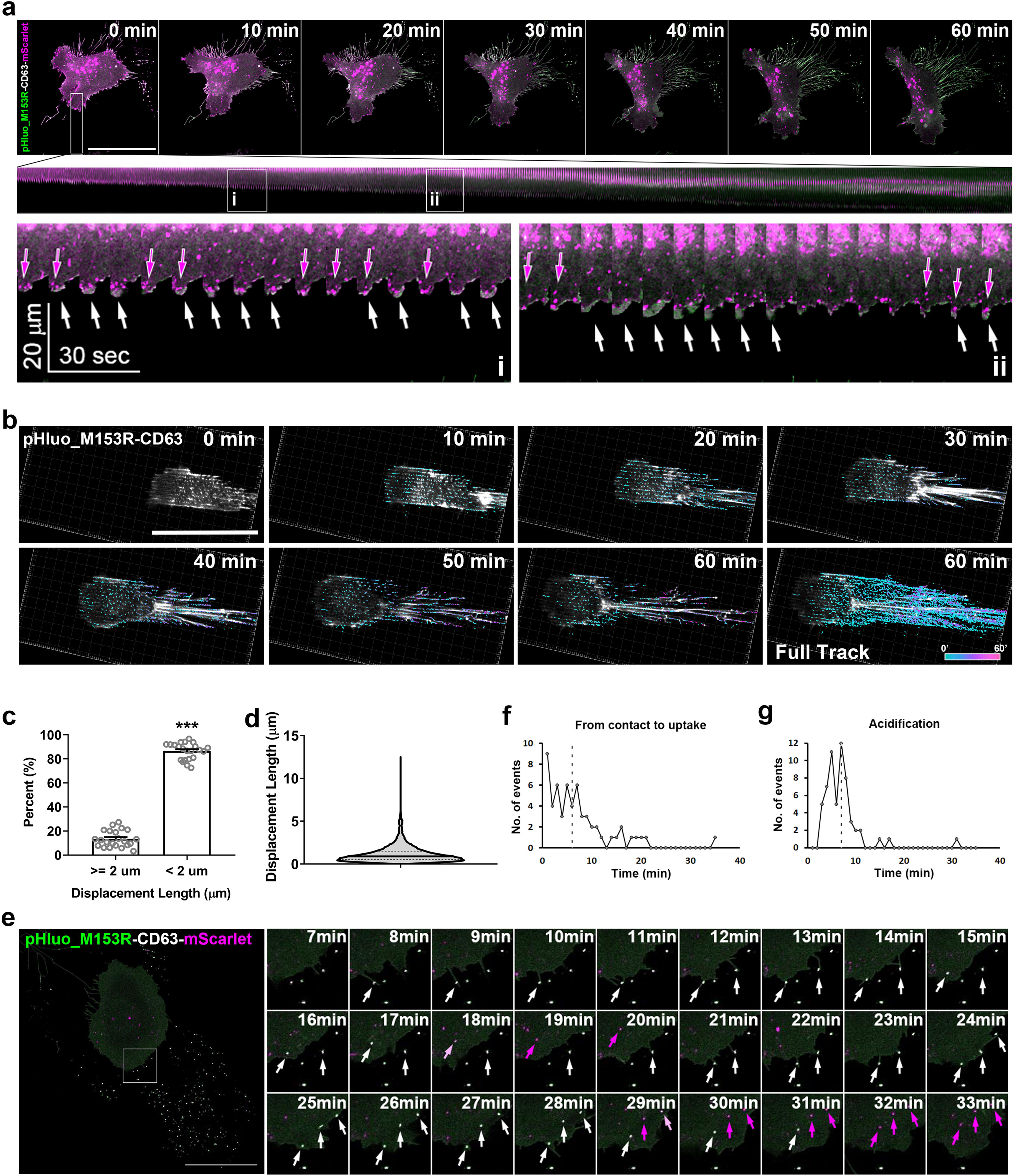
A dual reporter reveals MVB transport before fusion and endosome acidification after uptake. Live confocal microscopy was used to image pHluorin_M153R-CD63-mScarlet-expressing HT1080 cells. (**a**) Top: Images were captured at 10 sec intervals for 60 min (see Supplementary Video 4). Time-series images with 10 min intervals show frames at the beginning and end of the movie, with the rectangle across the leading edge in the 0 min frame indicating location from which the kymograph (middle) was derived. Scale bar, 50 µm. Middle: Kymograph shows events over time across the indicated rectangle. Two regions (i and ii) selected from kymograph (middle) are enlarged at the bottom of the panel. Magenta arrows in kymograph zooms indicate CD63-mScarlet-positive MVB trafficking and docking to protrusions while white arrows indicate MVB fusion and exosome secretion. (**b**) Time series from Supplementary Video 5, showing trajectories of tracked exosome deposits under the cell on a nanopatterned dish. Colors on heat map represent the time elapsed since deposition occurred. Note that cyan color shows newly deposited exosomes. Note the image at the right bottom corner (Full Track) shows the overlap of all of the tracks from the movie. n=23 cells from 4 independent experiments. Scale bar, 50 µm. (**c**) and (**d**) Analysis of mobility of exosome deposits from movies as in (**b**) with scatter dot plots in (**c**) (mean +/-standard error) showing the percent of exosome deposits/cell with displacement length > or < 2 µm, where each dot represents the median displacement length of exosome deposits in that category from each single cell. \*\*\**P*<0.001 (**d**) Total events (n=34052 from 23 cells) of displacement length are shown in a violin plot (median with interquartile range) from (**b**). (**e**) Time series images from live confocal microscopy of pHluorin_M153R-CD63-mScarlet-expressing HT1080 cells with 1 min intervals from Supplementary Video 6. White arrows indicate examples where filopodia contact exosome deposits. Magenta arrows indicate acidified exosome-containing endosomal compartments after exosome uptake. n=62 exosome uptake events from 10 cells from 5 independent experiments. Scale bar, 25 µm. (**f**) Quantitation of internalization time of exosome deposits after initial filopodial contact from (**e**). Median time from contact to uptake is marked by a dotted line. (**g**) Quantitation of time from exosome endocytosis to acidification as shown in (**e**). Median time of acidification is marked by a dotted line.

Both the trafficking of pHluo_M153R-CD63-mScarlet to cellular protrusions and the identified role of exosomes in directional migration ^20, 29-31^ suggest that exosomes should be secreted from the front of migrating cells. However, it has been difficult under standard conditions to definitively determine whether exosomes are secreted at the front or back (or both) of migrating cells, especially since CD63 localizes to both the plasma membrane and exosomes. To more definitively answer this question, we seeded pHluo_M153R-CD63-expressing HT1080 cells on a dish with a polymeric nanopatterned surface topography and performed confocal microscopy of moving cells (Fig. 4b, Supplementary Video 5). Due to the patterning of the substrate, new pHluo_M153R-CD63 deposits were easy to observe and track. From these movies we observed that new exosome deposits (depicted with cyan color in the heat map) clearly occurred at the front of the cell and were largely stationary as the cell moved over them, with reorganization of the deposits at the rear of the cell by retraction fibers.

Using a criteria of < 2 µm of displacement length during the movie as immotile, we find that 85% of deposits are immotile (Fig. 4c and d). While we cannot rule out that secretion also occurs from the rear of the cell (this is difficult to assess due the accumulation of fluorescence at the rear of the moving cell, presumably from ongoing secretion), some secretion clearly occurs at the front of the cell and thus could mediate the cellular directional sensing that has been shown to be a critical function of exosome secretion^20, 29, 30^.

It was previously shown by DIC and wide-field epifluorescence microscopy that exosomes can be captured by filopodia and endocytosed into cells^33^. Using the dual-color reporter, we observed a similar event in which migrating cells detect exosome deposits by touching them with filopodia, then migrate toward the exosome deposits and engulf them (Fig. 4e, white arrows, Supplementary Video. 6).

Although these experiments were not designed to identify mechanisms of endocytosis^34^, in some cases we observed micropinocytosis-like uptake by engulfment (25 min frame, top arrow). Of note, endocytosed exosomes turned from yellow (pseudocolored white) to red (pseudocolored magenta) over time, consistent with maturation and acidification of endosomes (Fig. 4e, magenta arrows). From the time-lapse images, we quantified the kinetics of internalization and acidification of endocytosed exosome deposits. The internalization time of exosome deposits from the initial filopodial contact was highly variable and ranged from 1 min to 20 min with a median internalization time of 6 min (Fig. 4f). The distance between the exosome deposits and the plasma membrane likely affects the length of time taken for exosome uptake after initial contact. However, acidification of endocytosed exosome deposits, as visualized by loss of the pHluorin signal, occurred in a more narrow time range, from 2 to 11 min with a median of 7 min (Fig. 4g).

## Discussion

Dynamic monitoring of EVs is critical to investigate roles of EV secretion in cell behaviors, including cell-cell communication and cell migration. Although we previously developed a pH-sensitive exosome secretion reporter, pHluorin-CD63^20^, the reporter was dim and could not be stably expressed in cells. By making a single amino acid mutation, we developed a stable and bright pH-sensitive reporter, pHluorin_M153R-CD63, for live imaging of MVB fusion and cellular interactions with extracellular exosomes. Using this reporter, we observed that exosomes are secreted at the front of migrating cells and left behind in exosome trails. We also observed strong pathfinding behavior of migrating cells along trails of extracellular exosomes. Finally, we made a dual color reporter that allows observation of not only exosome secretion but also internal trafficking events in both donor and recipient cells.

Recently, there has been much interest in observing EV release and interchange between cells. GFP-CD63 and fluorophores tagged to palmitoylation motifs have been used to observe uptake by recipient cells within tumors^35^. In zebrafish, recent papers used both pHluorin-CD63 (injected as plasmid DNA for transient expression) and a cyanine-based membrane probe to observe EV dynamics in the blood circulation ^36, 37^. Our new reporters provide important additions to these tools by allowing stable expression in diverse cell types and potentially in transgenic animals while allowing high resolution imaging of subcellular events.

A key question addressed in this study is whether pHluo_M153R-CD63-positive extracellular puncta correspond to exosomes. As shown by Western blot analysis and CLEM, pHluo_M153R-CD63 labels small but not large EVs. The extracellular puncta were not labeled with the plasma membrane marker, mCherry-CaaX, suggesting that they did not derive from the plasma membrane but rather from internal MVB. Consistent with that hypothesis, KD of the MVB docking factor Rab27a^23^ greatly reduced both the secretion of small EVs and the deposition of pHluo_M153R-CD63-positive extracellular puncta compared to control cells. Of note, there was a larger reduction in the area and intensity of extracellular CD63-positive deposits (Fig 2c-e) than in the number of secreted small EVs measured by NTA (Fig 2b). This discrepancy indicates the heterogeneity of small EVs and suggests the possibility of the release of small CD63-negative ectosomes from the plasma membrane^26^, as these would not be subject to regulation by Rab27a-mediated MVB docking. We also observed through immunofluorescent staining that pHluo_M153R-CD63-positive puncta are more frequently positive for Alix, an ESCRT accessory protein, than for TSG101, an ESCRT-I protein. These data are consistent with a previous finding that syntenin regulates the budding of CD63-positive ILVs into MVBs by interaction with Alix^10^, although TSG101 was also implicated in that process.

Exosome secretion promotes chemotaxis of several cell types^29-31^. Previously, we proposed a model in which exosomes secreted from cancer cells promote cancer cell chemotaxis in both an autocrine manner by secretion at the leading edge as well as in a paracrine manner by leaving behind exosome trails^29^. More recently, the Parent group reported that exosome trails released from migrating Dictyostelia induce streaming behavior ^31^. In this study, we observed that cancer cells migrate toward and over exosome deposits to use them as migration tracks in 2D and 3D tissue culture environments. Although it is not clear how much of this behavior is due to chemicals released from the exosomes to induce chemotaxis and how much is providing an adhesive^20^ and signaling migration track, through direct interaction, the overall behavior is clear. Future studies using our reporter, both *in vivo* and *in vitro*, in conjunction with molecular manipulations should be able to dissect further these behaviors.

Another striking finding, observed with our dual color reporter, is that cancer cells migrate toward exosome deposits and actively endocytose them. Heusermann *et al*. previously reported that exosomes enter cells through filopodia^33^, a similar behavior to that of viruses^38^. However, for the first time, we visualized and quantitated both exosome uptake and the subsequent acidification of the exosome-containing endosomal compartments. We anticipate that future studies may be able to use this dual-color reporter to monitor exosome interchange between cells and whether endocytosed exosomes are recycled and re-secreted. Furthermore, by combining with inhibitors, molecular interventions, and/or other reporter systems, the dual reporter may be useful for studying the endocytic fates of exosomes in diverse cell types.

Previous studies have reported that exosomes are likely secreted from the uropod or tail of migrating cells^39, 40^; however, those studies were performed with Dictyostelia and leukocytes that primarily use amoeboid locomotion for movement, involving the formation of posterior uropods. By contrast, mesenchymal migration and directional sensing of cells undergoing chemotaxis would seem to require exosome secretion at the leading edge of cells, at least for single cell autocrine migration events. Due to the fact that cells migrate over exosome paths, it has been difficult to discern where exosomes are secreted at a subcellular level. Using polymeric nanopatterned dishes, we observed clear deposition of pHluo_M153R-CD63-positive puncta at the front of migrating HT1080 cells. The deposited puncta stayed stationary as the cells moved over them and were left behind the migrating cells, with a small amount of reorganization along the way. We also observed that migrating cells left bright retraction fibers at the trailing edge attached to the exosome deposits. The strong adherence of the cells to the secreted exosomes in both the nanopatterned substrate experiments and in our other movies is consistent with the adhesive function of exosomes that we previously reported^20, 29^. Due to the bright accumulation of previously deposited exosomes, we could not observe or rule out additional exosome deposition at the rear of cells migrating on the nanopatterned plates; however, it is clear that exosomes are secreted at the cell front during migration.

To conclude, pHluo_M153R-CD63 is a stable and bright live cell reporter of exosome secretion. Migrating cells leave pHluo_M153R-CD63-positive exosome trails behind them and the exosome trails attract and promote migration of follower cells. With our dual color reporter pHluo_M153R-CD63-mScarlet, it is also possible to monitor MVB trafficking before fusion as well as exosome endocytosis. We anticipate that these reporters will be broadly useful to investigate regulation and functions of exosome secretion and uptake in diverse physiological conditions.

## Methods

### Cell culture and reagents

HT1080 fibrosarcoma cells were maintained in DMEM supplemented with 10% bovine growth serum (BGS). MDA-MB-231 breast cancer cells and B16F1 mouse melanoma cells were maintained in DMEM supplemented with 10% fetal bovine serum (FBS). HNSCC61 head and neck squamous cell carcinoma cells were maintained in DMEM supplemented with 20% FBS and 0.4 µg ml^− 1^µg/ml hydrocortisone. A lentiviral shRNA expression system, pLKO.1, was used to knockdown Rab27a (TRCN0000005296 (5’-CCGG-CGGATCAGTTAAGTGAAGAAA-CTCGAG-TTTCTTCACTTAACTGATCCG-TTTTT-3’) and TRCN0000005297 (5’-CCGG-GCTGCCAATGGGACAAACATA-CTCGAG-TATGTTTGTCCCATTGGCAGC-TTTTT-3’), Thermo Fisher Scientific) or scrambled control (5’-CCTAAGGTTAAGTCGCCCTCG-3’, Addgene Plasmid #26701). Viral particles were produced from 293FT cells by co-transfection with viral vectors. Cells were transduced by viral particles and selected using selection markers. For stable fluorescence expression, low-expressing cells were sorted by FACSAria III (BD Biosciences) and used for experiments. HT1080 cells expressing both pHluo_M153R-CD63 and mCherry-CaaX were sorted for red fluorescence by FACSAria III after transducing mCherry-CAAX in HT1080-pHluo_M153R-CD63. Primary antibodies were: anti-CD63 (ab68418, abcam, 1:500 for WB), anti-GFP (A11122, Invitrogen, 1:5,000 for WB), anti-TSG101 (ab30871, abcam, 1:1,000 for WB and 1:100 for IF), anti-flotillin (610820, BD Biosciences, 1:1,000 for WB), anti-GM130 (610822, BD Biosciences, 1:250 for WB), anti-Rab27a (69295, Cell Signaling, 1:1,000 for WB), anti-Alix (2171, Cell Signaling, 1:200 for IF) and anti-β-actin (Ac-74, Sigma, 1:5,000). HRP-, or Alexa Fluor 546-secondary antibodies were from Santa Cruz Biotechnology or Invitrogen, respectively. Pierce ECL Western Blotting Substrate and SuperSignal WestFemto Maximum Sensitivity Substrate (32106 and 34095, respectively, Thermo Fisher Scientific) were used as substrates for Western blotting. Blots were imaged using an Amersham Imager 600 (GE Healthcare Life Sciences) and analyzed using Fiji.

### Site-directed mutagenesis and cloning

Methionine153 on pHluorin in pcDNA3.1-pHluorin-CD63^20^ was mutated to Arginine using QuickChange II XL Site-Directed Mutagenesis Kit (Agilent) with a pair of primers (Forward, 5’-ACG AGC ACT TGG TGT ACA TCC GGG CAG ACA AAC AAA AGA ATG-3’ and Reverse, 5’-CAT TCT TTT GTT TGT CTG CCC GGA TGT ACA CCA AGT GCT CGT-3’). pcDNA3.1-pHluorin_M153R-CD63 was subcloned into pENTR/D-TOPO and then cloned into pLenti6/V5-DEST plasmid using Gateway recombination cloning (Thermo Fisher Scientific). mCherry-CaaX from pME-mCherry-CaaX (a generous gift from Chi-Bin Chien, University of Utah) was subcloned into pENTR/D-TOPO and then cloned into pLenti6/V5-DEST plasmid using Gateway recombination cloning (Thermo Fisher Scientific). mScarlet from pCytERM-mScarlet_N1 (a gift from Dorus Gadella, Addgene plasmid #85066) was cloned to the C-terminus of pHluo_M153R-CD63 in pLenti6/V5-DEST using Gibson Assembly Master Mix (NEB) with two pairs of primers (Forward, 5’-ACG AGC TGT ACA AGG GAT CCT AGA AGG GTG GGC GCG C-3’ and Reverse, 5’-CCC TTG CTC ACC ATG AAT TCC ATC ACC TCG TAG CCA CTT CTG A-3’ for pLenti6/V5-DEST-pHluo_M153R-CD63 and Forward, 5’-GAA GTG GCT ACG AGG TGA TGG AAT TCA TGG TGA GCA AGG GCG-3’ and Reverse, 5’-TCG GCG CGC CCA CCC TTC TAG GAT CCC TTG TAC AGC TCG T-3’ for pCytERM-mScarlet_N1). Site-directed mutagenesis and all subclonings were confirmed by sequencing (Genewiz).

### Isolation of EVs

To collect conditioned media, 80% confluent cells were cultured for 48 h in Opti-MEM (Thermo Fisher Scientific). Exosomes were isolated from conditioned media by serial centrifugation at 300 × g for 10 min, 2000 × g (4,000 rpm in Ti45 rotor) for 30 min, 10,000 × g for 30min (9300 rpm in Ti45), and 100,000 × g (30,000 rpm in Ti45) overnight to respectively sediment live cells, dead cells, debris and large EVs, and small EVs. Pellets of large and small EVs were resuspended in PBS and spun again in same conditions. Each pellet was resuspended in PBS and used for NTA using ZetaView (Particle Metrix) or for Western blot analysis. At the time of conditioned media collection, cells were trypsinized and counted to allow for estimation of the exosome secretion rate as number of exosomes divided by the number of cells and by the number of hours of media collection (48 h).

### Immunofluorescence staining

Cells on coverslips coated with FN (1 µg ml^−1^ overnight at 4°C) were permeabilized with 0.2% Triton X-100 in PBS after fixation with 4% paraformaldehyde in PBS. After blocking with 5% BSA in PBS, primary and secondary antibodies listed above were sequentially incubated the cells with PBS washes in between. After mounting coverslips on glass slides, Z-stack images were acquired with an LSM 510 laser scanning confocal microscope (CarlZeiss) equipped with a 63x/1.40 NA Plan Apo oil objective lens and processed by maximum intensity projection.

### Correlative light-electron microscopy

Cells were plated on high Grid-500 glass-bottom µ-Dishes (ibidi) coated with FN (1 µg ml^−1^) and cultured at 37 °C incubator with 5% CO_2_ for 1 day. Cells were initially washed in 0.1M sodium cacodylate buffer then briefly fixed in 2% paraformaldehyde. pHluo_M153R-CD63-overexpressing cells were identified using a Nikon Plan Apo 60x/1.40 oil immersion lens in a Nikon Eclipse TE2000E microscope equipped with a cooled charge-coupled device (CCD) camera (Hamamatsu ORCA-ER). The cells were then fixed in 2.5% glutaraldehyde in 0.1M cacodylate buffer, pH7.4 at room temperature (RT) for 1 hour then transferred to 4ºC, overnight. The samples were again washed in 0.1M cacodylate buffer, then incubated 1 hour in 1% osmium tetraoxide at RT, rinsed in 0.1M cacodylate buffer. Subsequently, the samples were dehydrated through a graded ethanol series, 3 exchanges of 100% ethanol, and then the glass-bottoms were taken off from the dishes, followed by critical point drying with a samdri-PVT-3D Tousimis critical point dryer. The dried coverslips were then mounted on an aluminum stub with a carbon adhesive tab and sputter coated with gold using the Cresssington Sputter Coater 108. After coating for 60 seconds, the samples were imaged using a Quanta 250ESEM.

### Live imaging in 2D and 3D

Cells were plated on glass-bottom MatTek dishes coated with fibronectin (1 µg ml^−1^) and maintained in complete media in a 37 °C incubator with 5% CO_2_. The next day, the media was changed to Leibovitz’s L-15 (Gibco)/10% serum or FluoroBrite DMEM (Gibco)/10% serum depending on the CO_2_ microscope environment. Live imaging of mCherry-CaaX/pHluo_M153R-CD63-expressing HT1080 cells was performed with a Nikon Eclipse TE2000E epifluorescence microscope equipped with a 37 °C chamber and a cooled CCD camera using a Nikon Plan Fluor oil 40x/1.30 or Super Fluor 20x/0.75 NA objective lens. Still images of pHluo_M153R-CD63 were captured from diverse live cells using a Nikon Plan Fluor oil 40x/1.30 lens on the Nikon Eclipse TE2000E. Time lapse movies of pHluo_M153R-CD63-expressing cells on nanopatterned glass-bottom dishes (800 nm width of both ridge and groove and 600 nm depth, Nanosurface Biomedical) coated with fibronectin (1 µg ml^−1^), and of cells expressing the dual reporter pHluo_M153R-CD63-mScarlet, were acquired with a Nikon A1R confocal microscope equipped with a Tokai Hit Incubation Chamber (37 °C with 5% CO_2_) using a Plan Apo 40x/1.3 NA oil immersion lens.

### Intravital imaging

#### Mouse intravital imaging

All mouse imaging and surgical protocols were approved by the University of Wisconsin Institutional Animal Care and Use Committee (IACUC) and were carried out in accordance with IACUC guidelines. For intravital experiments, 1×10^6^ MDA-MB-231 breast cancer cells expressing pHluorin_M153R-CD63-mScarlet were injected into the mammary fat pads of 8 week old female NOD/SCID mice. Imaging of mammary tumors was accomplished through surgical implantation of a mammary imaging window (MIW). Tumors were palpable at one week post-injection and MIWs were implanted at this time. Multiphoton microscopy of the pHluorin tumors commenced 2 days post MIW implant and was conducted on an Ultima IV (Bruker Nano Surfaces, Middleton, WI) using a Coherent Chameleon laser and Hamamatsu multi-alkali photomultiplier detectors. Data acquisition and scanning control was provided by PrairieView (Bruker Nano Surfaces, Middleton, WI). Mice were anesthetized using isoflurane, provided subcutaneous PBS hydration, and placed in a warming chamber mounted to the microscope stage for imaging. Respiration rates were monitored during the imaging period. Images were collected using a Nikon 40x 1.15 NA water immersion objective lens. Imaging of collagen and cells was performed simultaneously using a Chroma SHG/eGFP (430/40, 520/40) custom filter cube and 890 nm excitation.

#### Chick embryo intravital imaging

Fertilized chicken embryos were placed in a 371⍰°C incubator at 55% relative humidity on day 1 post-fertilization. Eggs were decanted into sterilized weigh boats on day three for *ex ovo* chorioallantoic membrane development. On day 10, 1 × 10^5^ HT1080 fibrosarcoma cells expressing pHluo_M153R-CD63/ mCherry-CaaX were injected (100 μL) in the direction of blood flow into the allantoic vein. 24 hours post injection, the *ex ovo* chicks were transferred to an intravital imaging chamber and placed under a temperature-controlled (371⍰°C) microscope. Images of HT1080 cells in the chick chorioallantoic membrane were acquired using an Olympus BX61 equipped with Olympus 20x 0.70 NA UApo/340 water immersion objective lens and a CCD camera (Hamamatsu ORCA-ER C4742-80-12AG) controlled with Volocity image acquisition software (PerkinElmer).

### Kymograph analysis

Cells expressing pHluo_M153R-CD63-mScarlet were plated on glass-bottom MatTek dishes coated with fibronectin (1 µg ml^−1^) and time lapse movies were acquired every 10 seconds with an A1R-HD25 confocal microscope equipped with a Tokai Hit Incubation Chamber (37 °C with 5% CO_2_) using a Plan Apo 40x/1.3 NA oil immersion lens. Region of interest was selected by a rectangular selection tool and a time-series montage was made by using Fiji (Image/Stacks/Make Montage).

### Image Analyses

#### Analysis of extracellular exosome deposits

Cell bodies were carefully selected and deleted from each image. The remaining pHluo_M153R-CD63 deposits were segmented from the background by thresholding and then measured for area and integrated intensity using Fiji (Analyze tab/Measure). For colocalization of pHluo_M153R-CD63 with ESCRT proteins, fluorescence signals surrounding cells were segmented from the background by thresholding and measured for colocalization of pHluo_M153R-CD63 with TSG101 or Alix using MetaMorph (Molecular Devices, Apps/Measure Colocalization).

#### Pathfinding Index

Cell migration trajectories were created using Fiji (Plugins/Tracking/Manual Tracking). Then, the degree of angle (°) was measured between each trajectory and the closest exosome trail using the Angle Tool in Fiji. Degree was converted into radian (θ) and cosine value of radian was calculated using Excel (Microsoft), e.g. cos 0° = 1, cos 90° = 0, and cos 180° = −1. Only single migratory cells were selected for the analysis.

#### Nanopatterned substrates

To track and analyze mobility of pHluo_M153R-CD63 deposits on nanopatterned dishes, Imaris (Bitplane, vesicle tracking algorithm) and Fiji (Plugins/Mosaic/Particle Tracker 2D/3D) were used. Immobile deposits were defined as deposits with displacement length < 2 µm during one hour.

### Numbers and Statistics

For both quantitated data and representative images from experiments, the n values and independent experiment numbers are listed in the figure legends. Quantitated imaging data were acquired from at least 3 independent experiments and multiple images or movies on multiple cells were captured. Cell numbers to be quantitated for each experiment were determined by our experience and data were excluded only if there was an obvious reason for poor data, such as dead or sick-looking cells. For non-quantitated Western blots (e.g. checking knockdown), they were generally performed a single time. All datasets were tested for normality using the Kolmogorov-Smirnov normality test in GraphPad Prism. Non-parametric data groups were compared by the Mann-Whitney test and plotted as scatter plots with median and interquartile. Parametric data were compared using Student *t*-test and plotted as mean+/-standard error in scatter plots. The violin plot with median and interquartile range was created by Graphpad Prism to show all data.

## Acknowledgements

This study was funded by grants R01GM117916 and R01CA206458 to A.M. Weaver and 3U19CA179514-05S1 to R.J. Coffey, R01CA216248 to S.M. Ponik, and supported by a voucher from Vanderbilt CTSA grant UL1 RR024975. We thank Ian Macara for use of his Nikon A1R confocal microscope and Matt Tyska, as well as members of the Weaver laboratory for extensive discussion. The content of this article is solely the responsibility of the authors and does not necessarily represent the official views of the National Institutes of Health. Flow cytometry sorting was performed in the Vanderbilt University Medical Center (VUMV) Flow Cytometry Shared Resource. The VUMC Flow Cytometry Shared Resource is supported by the Vanderbilt Ingram Cancer Center (P30 CA68485) and the Vanderbilt Digestive Disease Research Center (DK058404). Data analysis using Imaris and scanning electron microscopy were performed in part through the use of the Vanderbilt Cell Imaging Shared Resource (supported by NIH grants CA68485, DK20593, DK58404, DK59637 and EY08126).

## Author Contributions

B.H.S. performed the experiments and data analyses and prepared the figures. A.v.L., J.G., D.I., and R.P. performed experiments. B.H.S., A.Z., S.P., and A.M.W designed the experiments. B.H.S. and A.M.W wrote the manuscript, with editing and input from coauthors.

