## Supplemental information for "pHluo_M153R-CD63, a bright, versatile live cell reporter of exosome secretion and uptake, reveals pathfinding behavior of migrating cells"

Sung et al.

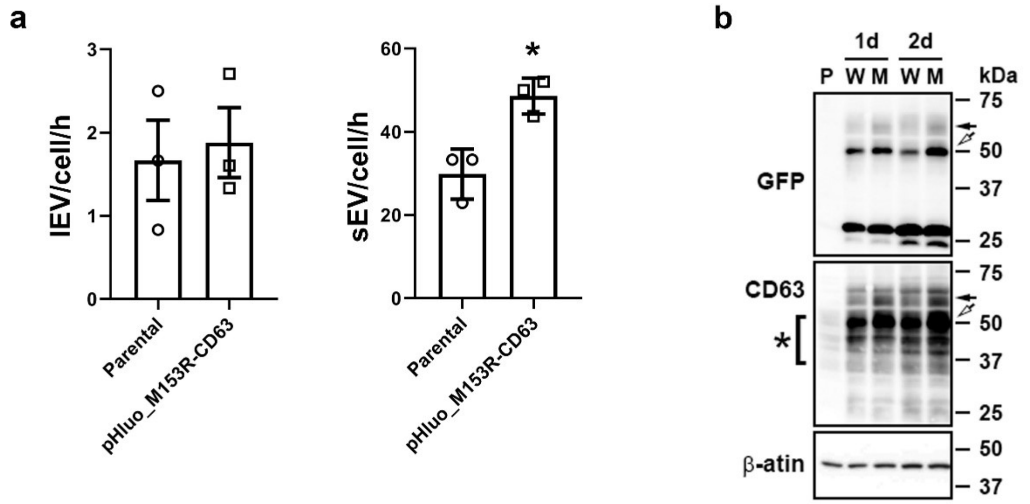


**Supplementary Figure 1. Overexpression of pHluorin_M153R-CD63 increases secretion rate of exosome-like small EVs.** (**a**) EV secretion rates measured by NTA. lEV, large EV. sEV, small EV. *$P$<0.05. (**b**) Western blots for transiently transfected HT1080 cells. P, parental cell lysate. W, pcDNA-pHluorin-CD63-transfected cell lysate. M, pcDNA-pHluorin_M153R-CD63-transfected cell lysate. 1d, 1 day after transfection. 2d, 2 days after transfection Black arrows indicate full length pHluorin_M153R-tagged CD63, which is shifted due to the GFP moiety of 27 kDa, while white arrows indicate potential cleaved form of CD63 tagged with pHluorin_M153R. Asterisk indicates cellular CD63, which has a broad range due to glycosylation.

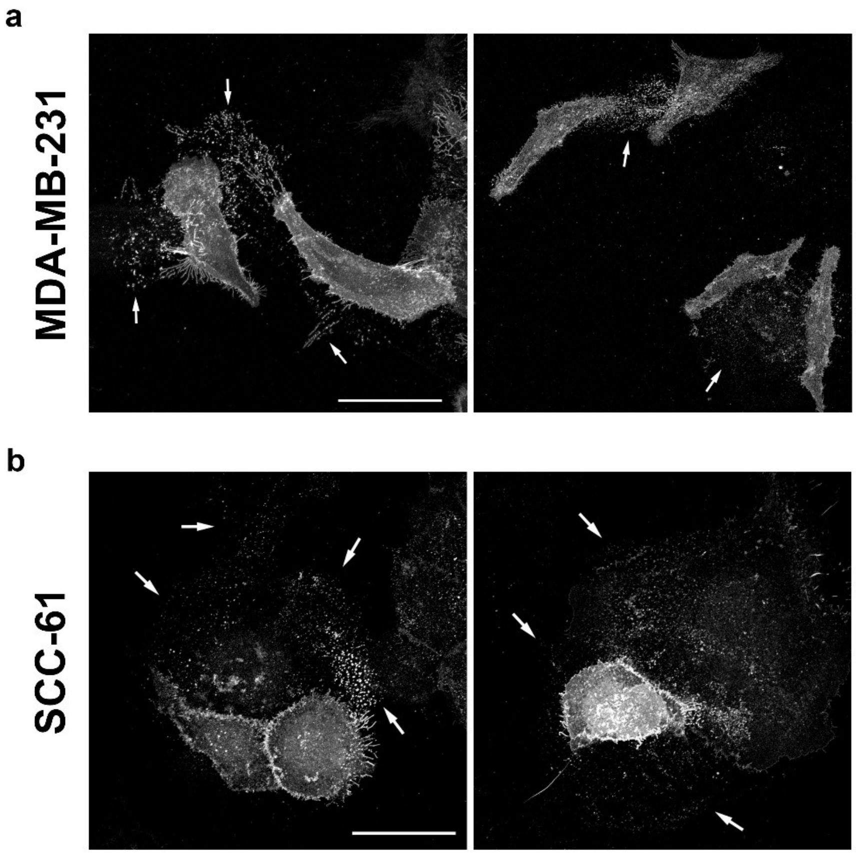


**Supplementary Figure 2. pHluorin_M153R-CD63-positive trails are deposited from diverse cancer cell types.** (**a**) Fluorescence images of live MDA-MB-231 human breast cancer cells stably expressing pHluo_M153R-CD63. (**b**) Fluorescence images of live HNSCC-61 human head and neck cancer cells stably expressing pHluo_M153R-CD63.

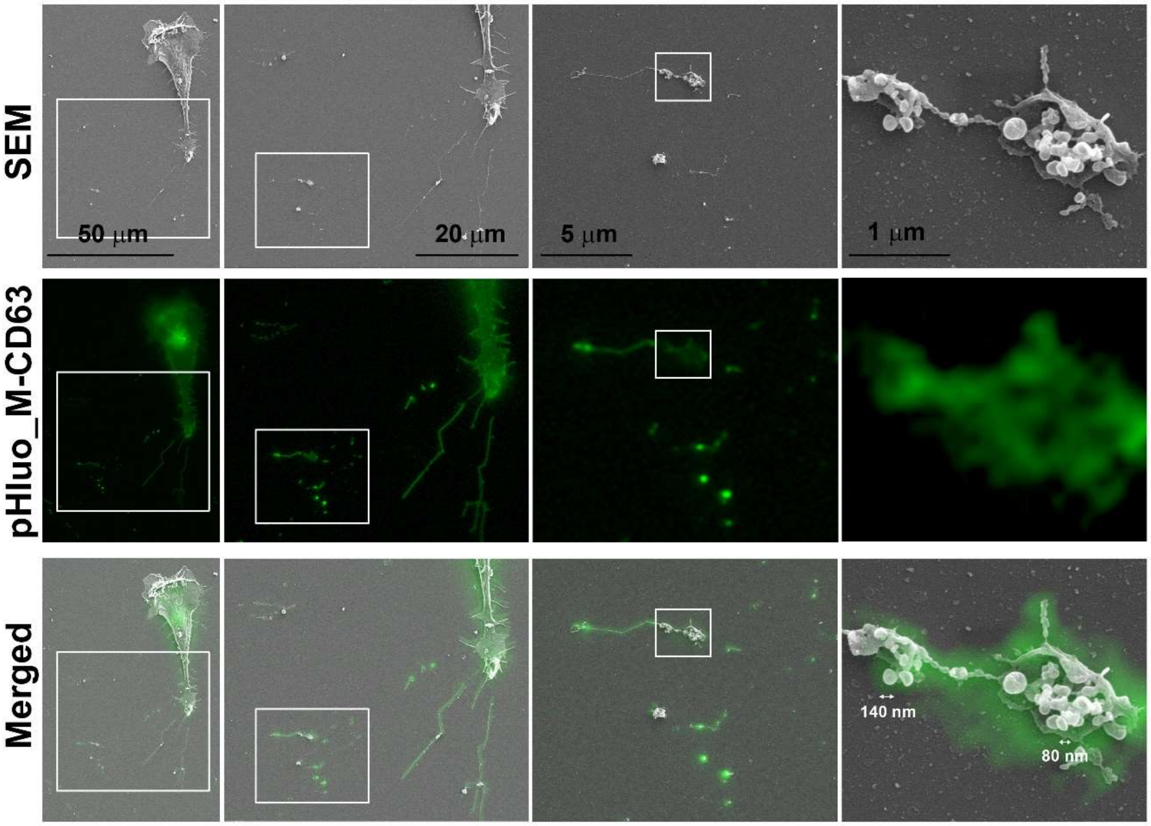


**Supplementary Figure 3. pHluorin_M153R-CD63-positive trails mark exosome deposits**. Correlative light-electron microscopy of pHluo_M153R-CD63-stably expressing HT1080. Zoom-in images from the left panel are shown to the right panel.

**Supplementary Video 1**. Migrating HT1080 cell stably expressing pHluo_M153R-CD63 and mCherry-CaaX (upper panel) and transiently expressing pHluo-CD63 (lower panel). Time-lapse images were taken on a fibronectin-coated MatTek dish (1 µg $ml^{-1}$) every 30 sec. Note the extracellular puncta left behind the migrating cell and the dimmer fluorescence from pHluo-CD63 compared with pHluo_M153R-CD63. Scale bars, 50 µm.

**Supplementary Video 2**. Migrating HT1080 cells in a 2D environment exhibit pathfinding behavior over pHluo_M153R-CD63 deposits. Time-lapse images were taken on a fibronectin-coated MatTek dish (1 µg $ml^{-1}$) every 5 min. Scale bar, 50 µm.

**Supplementary Video 3**. Migrating HT1080 cells in a 3D environment exhibit pathfinding behavior over pHluo_M153R-CD63 deposits. Time-lapse images were taken in collagen gels (1.5 mg $ml^{-1}$) every 10 min. Scale bar, 50 µm.

**Supplementary Video 4**. Live confocal imaging of pHluo_M153R-CD63-mScarlet reveals that MVBs are trafficked toward and exosomes are secreted at the leading edge of migrating cells. Time-lapse images were taken every 10 sec. The white rectangle across the leading edge is a region for kymograph shown in Fig. 4a.

**Supplementary Video 5**. Live confocal imaging of pHluo_M153R-CD63 on a polymeric nanopatterned dish. Time-lapse images were taken every minute. Note that pHluo_M153R-CD63 deposits at the retraction fibers stayed stationary as the cells moved over them.

**Supplementary Video 6**. Endocytosis and acidification of extracellular exosome deposits. Live confocal imaging of pHluo_M153R-CD63-mScarlet with frames taken every minute. Note that the HT1080 cell migrates toward the exosome deposits and contacts them using filopodia. Contacted exosome deposits (white arrows) are endocytosed and acidified in endosomal compartments (magenta arrows).
